# Towards Spider Glue: Long-read scaffolding for extreme length and repetitious silk family genes AgSp1 and AgSp2 with insights into functional adaptation

**DOI:** 10.1101/492025

**Authors:** Sarah D. Stellwagen, Rebecca L. Renberg

**Affiliations:** Department of Biological Sciences, University of Maryland Baltimore County, Baltimore, MD 21250; General Technical Services, Adelphi, MD 20783

**Keywords:** spidroin, aggregate glue, long reads, full-length gene, spider silk, *Argiope trifasciata*

## Abstract

The aggregate gland glycoprotein glue coating the prey-capture threads of orb weaving and cobweb weaving spider webs is comprised of silk protein spidroins (spider fibroins) encoded by two members of the silk gene family. It functions to retain prey that make contact with the web, but differs from solid silk fibers as it is a viscoelastic, amorphic, wet adhesive that is responsive to environmental conditions. Most spidroins are extremely large, highly repetitive genes that are impossible to sequence using only short-read technology. We sequenced for the first time the complete genomic Aggregate Spidroin 1 (AgSp1) and Aggregate Spidroin 2 (AgSp2) glue genes of *Argiope trifasciata* by using error-prone long reads to scaffold for high accuracy short reads. The massive coding sequences are 42,270 bp (AgSp1) and 20,526 bp (AgSp2) in length, the largest silk genes currently described. The majority of the predicted amino acid sequence of AgSp1 consists of two similar but distinct motifs that are repeated ~40 times each, while AgSp2 contains ~48 repetitions of an AgSp1-similar motif, interspersed by regions high in glutamine. Comparisons of AgSp repetitive motifs from orb web and cobweb spiders show regions of strict conservation followed by striking diversification. Glues from these two spider families have evolved contrasting material properties in adhesion, extensibility, and elasticity, which we link to mechanisms established for related silk genes in the same family. Full-length aggregate spidroin sequences from diverse species with differing material characteristics will provide insights for designing tunable bio-inspired adhesives for a variety of unique purposes.

## 1 Introduction

Spiders use a suite of remarkable silk and silk-derived materials for various applications during their life cycles, from wrapping prey and egg cases, to creating webs, lifelines, and prey capture glues. Aggregate gland glue is produced by spiders within the superfamily Araneoidea and functions to retain prey that get caught in a spider’s silken trap. This material is an adhesive glycoprotein (Tillinghast, 1981) that coats the capture spiral silk of orb web weavers, the silk trip lines of cobweb weavers, and the globule of glue at the end of the bolas of bolas spiders (Fig. 1). The glue is produced within a spider’s abdomen by two pairs of aggregate glands that terminate with valved spigots on the spinnerets (Sekiguchi, 1952).

**Figure 1:**
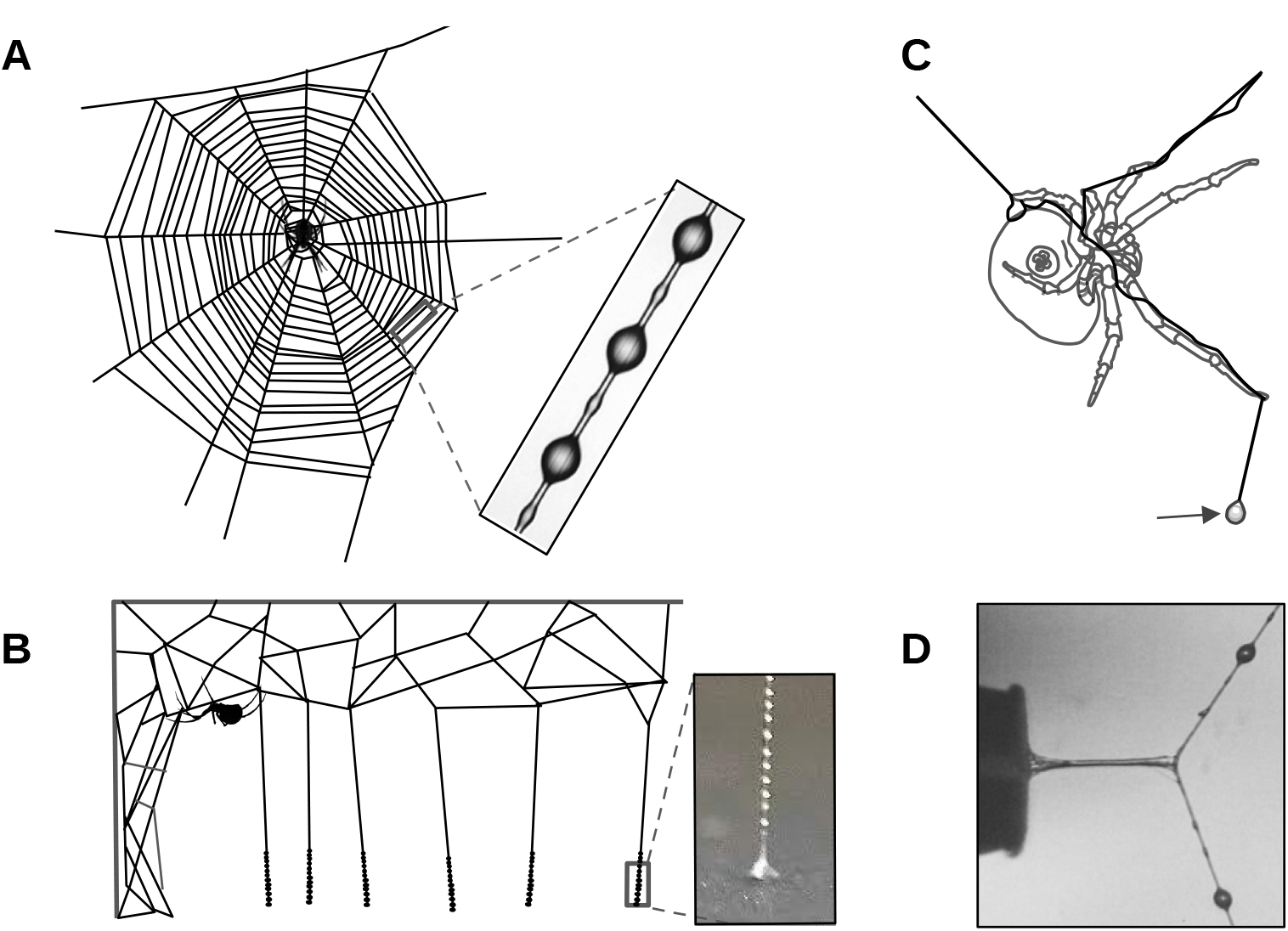
Aggregate spider glue from three spider types. (A) Orb weavers coat the web’s sticky capture spiral with glue, (B) the bolas spider creates a large droplet of glue specializedfor capturing moths, and (C) cobweb weavers cover the lower portion of their triplines with glue to create ‘gumfoot’ threads. (D) A stretching glue droplet after contacting a probe that was subsequently with drawn at a constant rate. Inset images courtesy of Brent Opell.

In orb weavers, each pair of semi-circle-shaped glue spigots surrounds a flagelliform silk spigot, which produces the supporting thread as the glue is concurrently extruded. The amorphous glue is deposited on the flagelliform threads as a continuous cylinder, however as the two coated threads merge to form a single strand, hygroscopic low molecular mass compounds (LMMC) and salts absorb atmospheric moisture. Surface tension then causes the cylinder of glue to separate into equally spaced droplets (Fig. 1A)(Plateau, 1873; Boys, 1889; Edmonds and Vollrath, 1992). A single droplet contains a core of adhesive glycoprotein (Opell and Hendricks, 2010; Sahni et al., 2010) surrounded by an outer aqueous layer containing the LMMC and salts that retain water and lubricate the core of glue (Fischer and Brander, 1960; Anderson and Tillinghast, 1980; Tillinghast and Christenson, 1984; Townley et al., 1991; Opell et al., 2011, 2013; Sahni et al., 2014).

Orb weaving spider glue acts as a viscoelastic solid, exhibiting viscous properties at more rapid extension rates, and elasticity at slow extension rates (Sahni et al., 2010). This allows the glue to adhere to fast-moving insects at interception, and retain insects while a spider travels to and subdues them. The stickiness of the glue droplets, however, is constrained by the force required to break the flagelliform support fibers, as glue that does not preemptively release results in damaged webs as an insect pulls free by breaking the support fibers (Agnarsson and Blackledge, 2009). Capture spiral threads also exhibit a suspension bridge mechanism, allowing forces to be distributed across multiple droplets and reducing thread peeling that would occur if only droplets on the ends of a contacted thread contributed to adhesion (Opell and Hendricks, 2007).

Orb weaver glue responds instantly to humidity, which changes droplet volume and influences the performance of the glue (Opell et al., 2011; Sahni et al., 2011a). Orb weavers produce glue optimized for their habitat’s humidity, for example the glue of species from drier habitats becomes over-lubricated when exposed to higher humidity, while glue from species occupying wetter habitats becomes too viscous in drier conditions. Moreover, temperature and ultraviolet exposure also influence performance optima (Stellwagen et al., 2014, 2015).

In contrast to orb weavers, cobweb weavers connect major ampullate silk threads from their webs to the ground, depositing aggregate glue on the lower portions to produce trip lines aimed at ambulatory prey (Fig. 1B). The trip lines, or ‘gumfoot’ threads, easily release from the ground, pulling insects into the air when accidentally intercepted, and the prey hang suspended until the spider can subdue them (Argintean et al., 2006). Unlike orb weaver glue, cobweb glue is considered a viscoelastic liquid, is resistant to changes in humidity, and is less extensible (Sahni et al., 2011a). Gumfoot droplets lack the glycoprotein core visible in those of orb weavers, and tend to flow together and coalesce rather than simply swelling when exposed to increased humidity (see figure 1 in Sahni et al. (2011a)). Cobweb aggregate gland pairs are morphologically distinct, and have likely diverged in function as well (Kovoor, 1977; Townley and Tillinghast, 2013; Clarke et al., 2017).

Bolas spiders capture male moths using a single large glue droplet that hangs at the end of a silk strand (Fig. 1C). By mimicking female moth pheromones, the bolas spider lures the males to within striking range, and then swings their sticky snare until contact is made (Eberhard, 1977). The remarkable glue of these spiders is able to adhere in spite of a moth’s scales, which easily detach allowing them to escape other forms of predation (Sahni et al., 2011b). The aggregate glue of related moth specialist *Cyrtarachne akirai* Tanikawa, 2013 was recently characterized and was shown to have unique properties that allow the glue to spread and penetrate a moth’s scales (Diaz et al., 2018).

The proteins that form spider silks and glues are from the same family termed spidroins (‘spider fibroins’) (Hinman and Lewis, 1992), and are often encoded by very long (>5kb), repetitive, single exon genes. Functional specialization of spidroin paralogs is associated with the evolutionary success of spiders (Shultz, 1987), particularly the central repetitive motifs that are responsible for the diversity of silk material properties (Gosline et al., 1999). The first full-length spidroin cDNAs were isolated from the orb weaving spider *Argiope bruennichi* (Scopoli, 1772) in 2006 (Zhao et al., 2006). The ~9 kb *A. bruennichi* cylindrical gland spidroins (CySp1 and CySp2; synonymous with tubiliform spidroins, TuSp), encode for silk that is used to wrap egg sacs. The gene that encodes for the most well known spidroin, major ampullate silk (more colloquially known as dragline silk), was fully sequenced from the black widow *Latrodectus hesperus* Chamberlin & Ivie (1935) genomic DNA the following year (Ayoub et al., 2007). To date, 19 large (>5kb), full-length, and classified spidroins (i.e., assigned to a specific silk gland source) have been described (Table 1). Several other smaller or unclassified silk gland genes have also been characterized (Babb et al., 2017).

**Table 1:** Full length, classified spidroins with >5 kb complete coding sequences (including start and stop codons), ordered from smallest to largest. Asterisk indicates a correction reported in Han and Nakagaki (2013) but not corrected on GenBank. CySp is synonymous with TuSp.

| Species | Spidroin | bp (cds) | Introns | Reference | Accession # |
| --- | --- | --- | --- | --- | --- |
| *Araneus ventricosus* | MiSp | 5298 | Yes | Chen et al. (2012) | JX513956.1 |
| *Araneus ventricosus* | TuSp1 | 5766 | No | Wen et al. (2017) | MF192838.1 |
| *Latrodectus hesperus* | MiSp_v1 | 6564 | No | Vienneau-Hathaway et al. (2017) | KX584003.1 |
| *Nephila clavipes* | MaSp-g | 7398 | Yes | Babb et al. (2017) | MWRG01015344.1 |
| *Nephila clavipes* | TuSp | 8631 | No | Babb et al. (2017) | MWRG01000785.1 |
| *Argiope bruennichi* | CySp1 | 8952 | No | Zhao et al. (2006) | AB242144.1 |
| *Latrodectus hesperus* | MaSp1 | 9390 | No | Ayoub et al. (2007) | EF595246.1 |
| *Argiope argentata* | TuSp1 | 9462 | No | Chaw et al. (2018) | MF962652.2 |
| *Nephila clavipes* | AcSp | 9636 | No | Babb et al. (2017) | MWRG01010975.1 |
| *Argiope bruennichi* | CySp2* | 9657 | No | Zhao et al. (2006) | AB242145.1 |
| *Nephila clavipes* | MaSp-h | 10014 | No | Babb et al. (2017) | MWRG01038219.1 |
| *Argiope bruennichi* | MaSp2 | 10083 | No | Zhang et al. (2013) | JX112872.1 |
| *Araneus ventricosus* | AcSp | 10338 | No | Wen et al. (2018) | MG021196.1 |
| *Nephila clavipes* | PiSp | 10554 | No | Babb et al. (2017) | MWRG01012117.1 |
| *Latrodectus hesperus* | MaSp2 | 11340 | No | Ayoub et al. (2007) | EF595245.1 |
| *Argiope argentata* | AcSp1 | 13440 | No | Chaw et al. (2014) | KJ206620.1 |
| *Argiope argentata* | PySp1 | 17280 | No | Chaw et al. (2017) | KY398016.1 |
| *Nephila clavipes* | AgSp-d | 17820 | No | Babb et al. (2017) | MWRG01016148.1 |
| *Latrodectus hesperus* | AcSp1 | 18999 | No | Ayoub et al. (2013) | JX978171.1 |

Initial evidence for aggregate glue gene sequences was derived from what were thought to be two full-length transcripts encoding the glue proteins of orb weaver *Nephila clavipes* (Linnaeus, 1767), and were named ASG1 (Aggregate Spider Glue; 1,221 bp) and ASG2 (2,145 bp) (Choresh et al., 2009). Several years later, it was determined that ASG1 expression is not unique to aggregate glands, and that the ASG2 sequence, while mostly correct, contained a short insertion (Collin et al., 2016). Collin et al. (2016) also discovered that the predicted ASG2 C-terminus aligns with other spidroins, and it was suggested ASG2 be renamed to AgSp1 (Aggregate Spidroin 1) to follow gene family convention, which we have adopted here. This study extended the known sequence to 2,821 bp, however the complete gene including the 5’ end encoding the predicted N-terminus still remained unidentified. During sequencing of the first orb weaving spider genome, which focused on describing spidroins from *N. clavipes*, the 5’ sequence for AgSp1 was discovered (Babb et al. (2017); called AgSp-c, following the study’s own spidroin naming scheme). While Babb et al. (2017) were unable to bridge a gap between the 5’ (7,660 bp) and 3’ (3,446 bp) ends of the gene, the known length of AgSp1 was extended to at least 11,106 bp. This study further reported three other aggregate gland spidroins (AgSp-a, AgSp-b, and AgSp-d).

We combined Illumina and Oxford Nanopore technologies to sequence AgSp1 from orb weaving spider *Argiope trifasciata* (Forskål, 1775), a large cosmopolitan species that is abundant in our location, and which has had many aspects of its biology extensively researched. During gene expression analyses to confirm high expression of AgSp1, we discovered and subsequently assembled another highly expressed spidroin, which we call AgSp2. Furthermore, we analyzed aggregate glue transcript data from three orb weavers and four cobweb weavers to discover and compare the repetitive motifs of AgSp1. While hygroscopicity is the result of the LMMC and salts, which provide tunable hydration and lubrication of the glycoprotein in orb weavers, less is known about what is responsible for differences in other functional properties of aggregate glues across species. It is not clear whether it is changes in the protein sequence, the glycosylations that are added to the protein during post-translational modification, or more likely a combination of these and other variables.

We hypothesize that described differences in material properties will be reflected in the predicted protein backbone of the glue of orb weaving and cobweb weaving spiders, as inferred from mechanisms established for related spidroins. Spider glue droplets exhibit 1) adhesion, typically to insect prey, 2) extensibility, to remain in contact with struggling prey, and 3) elasticity, to retain prey after extension. Glycosylations have been attributed to glue adhesion (Singla et al., 2018), however potential mechanisms for extensibility and elasticity have only been described for other silks. Orb web weaver glue droplets are more extensible (i.e. stretch farther, Fig. 1D) than cobweb weaver droplets (Sahni et al., 2011a), and we expect to find sequence similarity related to increased extensibility in paralogous genes from the spidroin family to be more prominent in orb weaver glue. Elasticity, or the ability of a material to resume its original shape, is an important property for spider glues. As prey struggle and glue droplets extend, glue elasticity causes droplets to return to their original shape, retaining prey the web. We expect AgSp1 to contain sequence that may contribute to the elastic properties of the predicted protein.

Understanding the underlying mechanisms that provide aggregate spider glue with its diverse properties will provide insight into designing novel bio-inspired adhesives. Synthetic spider glues would not require replicating the spinning process that transforms liquid protein dope into solid threads within a spider’s gland duct, traits that have made synthetic silks challenging to scale. The aggregate glue is extruded without the same processing (Townley and Tillinghast, 2013), and synthetic versions will provide water-soluble and biodegradable solutions to problems ranging from pest control to temporary sealants.

## 2 Results

We sequenced, assembled, and corrected two highly expressed spidroin genes from the aggregate glue glands of orb weaving spider species *Argiope trifasciata*. To achieve full-length sequences, we used high-accuracy RNAseq short reads from Illumina to correct error-prone gDNA long reads from Oxford Nanopore. Aggregate Spidroin 1 (AgSp1) is encoded by 42,270 bp from two similarly sized exons spanning ~48,960 bp of genomic DNA, which includes a single ~6,690 bp central intron (Fig. 2; accession: MK138561). AgSp2 is encoded by 20,526 bp from a short 5’ and large 3’ exon, and contains an ~31,455 bp intron, in total spanning ~51,981 bp of gDNA (Fig. 3, accession: MK138559). We collected RNAseq short reads for long read correction, resulting in highly accurate coding sequences for these two genes, however apart from a few reads from pre-mRNA, RNAseq reads do not cover genomic intron sequence and is therefore uncorrected. Long read scaffolds and pre-mRNA reads did allow identification of putative splice sites for both aggregate spidroins (Supplementary Fig. S4).

**Figure 2:**
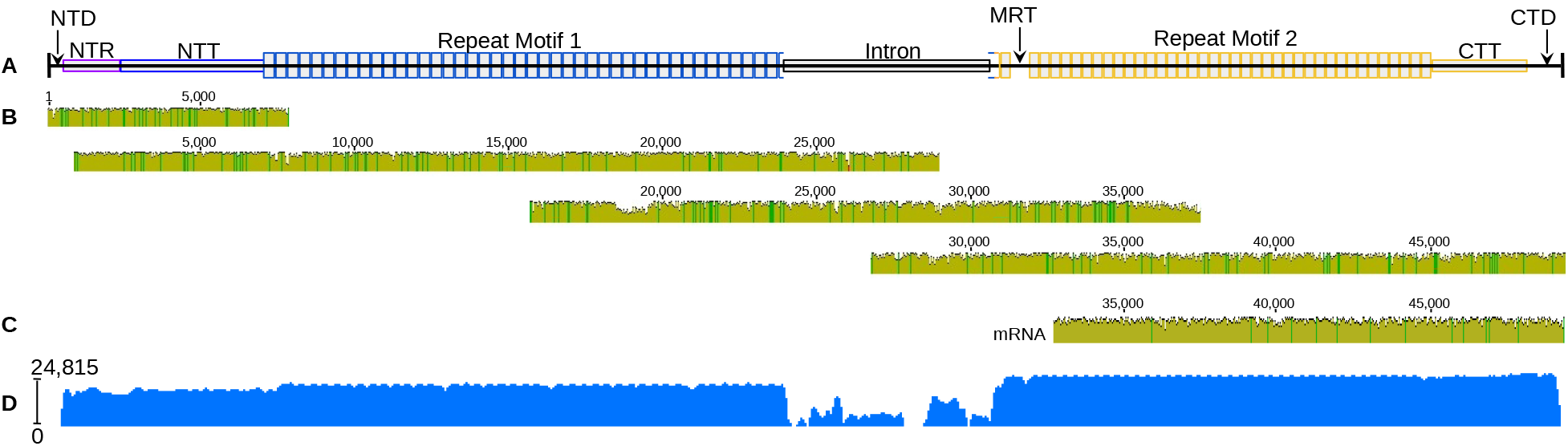
*A. trifasciata* Aggregate Spidroin 1 (AgSp1) schematic, aligned Oxford Nanopore gDNA reads, longest alignable mRNA read, and mapped read coverage. (A) AgSp1 consists of 42,270 bp of coding sequence and ~6,690 bp of intronic sequence, totalling ~48,960 bp of genomic sequence (intronic sequence could not be corrected with short reads derived from mRNA). Abbreviations correspond to regions of the predicted protein: NTD = N-terminal domain; NTR = N-terminal repeats; NTT = N-terminal transition; MRT = mid-rep eat transition; CTT = C-terminal transition, CTD = C-terminal domain. (B) Individual alignment of four Oxford Nanopore reads to the consensus AgSp1 together cover the entirety of the gene. (C) Alignment of a 16.4 kb mRNA transcript to the consensus AgSp1. (D) Log read coverage of Illumina RNAseq data generated from aggregate gland tissue mapped to AgSp1.

**Figure 3:**
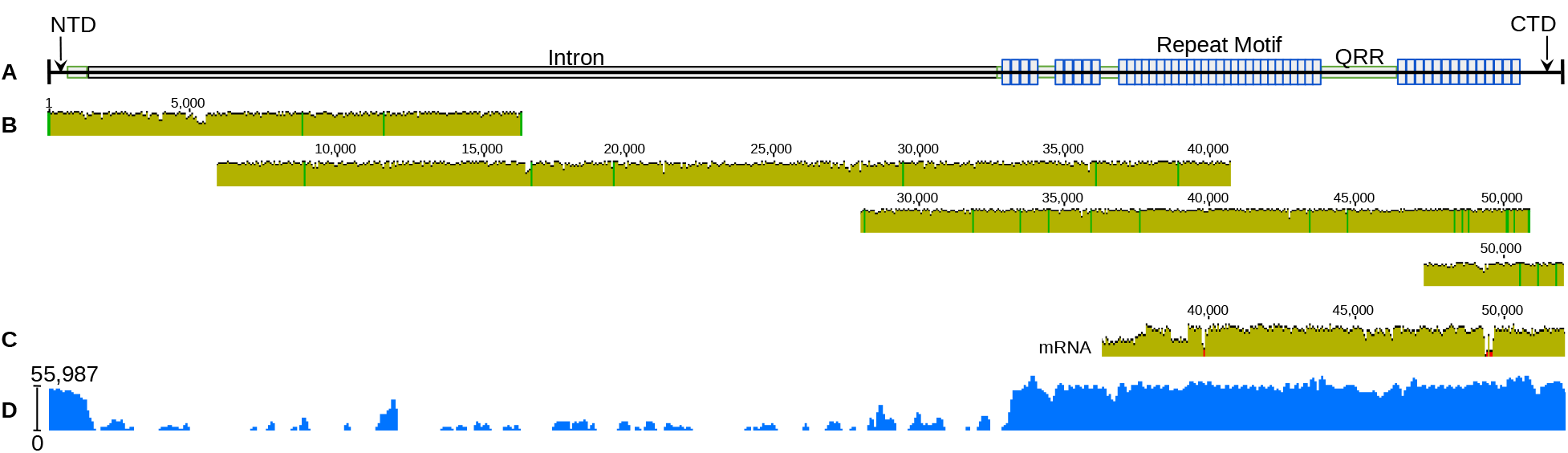
*A. trifasciata* Aggregate Spidroin 2 (AgSp2) schematic, aligned Oxford Nanopore gDNA reads, longest alignable mRNA read, and mapped read coverage. (A) AgSp2 consists of 20,526 bp of coding sequence and ~31,455 bp of intronic sequence, totalling ~51,981bp of genomic sequence (intronic sequence could not be corrected with short readt derived from mRNA). Abbreviations correcpond to regions of the predicted protein: NTD = N-terminal domain; QRR = glutamine -rich region (all regions in green); CTD = C-terminal domain. (B) Individual alignment of four Oxford Nanopore reads to the con-ensus AgSp2 together cover the entirety of the gene. (C) Alignment of a 16.5 kb RNA transcript to the consensus AgSp2. (D) Log read coverage of Illumina RNASeq data generated from aggregate gland tissue mapped to AgSp2.

AgSp1 and AgSp2 are members of the spidroin protein family (Collin et al., 2016) and are encoded by the same general N-terminus/Repeat(n)/C-terminus pattern found in this family, however the size and internal organization are distinct. We have broken the predicted protein of each spidroin into several regions. AgSp1 regions include: 1) an N-terminal domain (NTD), 2) a short region of N-terminal repeats (NTR), 3) an N-terminal transition (NTT, region with degenerate, repeat-similar structure), 4) 43 iterations of repeat motif 1 (RM1), 5) 38 iterations of repeat motif 2 (RM2), 6) a C-terminal transition (CTT, region with degenerate, repeat-similar structure), and 7) a C-terminal domain (CTD) (Fig. 2 and Supplementary Fig. S1). AgSp2 consists of an NTD and areas rich in glutamine (QRR - glutamine-rich region) that intersperse variable sized blocks of an iterated single repeat motif (RM) (Fig. 3 and Supplementary Fig. S2).

Current RNA sequencing technology does not provide reliable full-length sequencing of extreme-sized transcripts, however we were able to sequence a ~16.5 kb continuous RNA read from each aggregate spidroin using Oxford Nanopore’s MinlON and direct RNA sequencing kit (Figs. 2C and 3C). These enormous reads terminated with their 5’ ends still in the repetitive regions a few iterations before the genes’ respective introns. While most classified spidroins >5 kb in length are single exon (Table 1), AgSp1 and AgSp2 have large, single introns (Figs. 2 and 3).

### AgSp1

The 5’ end of AgSp1 encodes for the N-terminal domain (NTD; Spidroin_N domain superfamily BLASTx e-value 3.36e-14). Following the NTD, AgSp1 contains a short ~1,758 bp region that consists of translated TGSYITGESGSYD repetitions before connecting to the N-terminal transition encoding region. The N- and C- terminal transition regions that flank the iterative central repeat motif blocks (discussed below), consist of sequence that is unique enough to be assembled using only short reads, however the amino acid translation clearly resembles and aligns with the repeat motifs. These degenerate repeats become increasingly less conserved as the sequence approaches either terminus (Supplementary Fig. S1). The C-terminal transition region is shorter than the N-terminal transition, and maintains the same organization as the internal repeats, with the majority of variation occurring towards the end of each repeat iteration. The C-terminus has been previously described (Choresh et al., 2009; Collin et al., 2016).

The central repetitive region of AgSp1 contains at least two distinct repeat motifs. There are 43 iterations of repeat motif 1 (RM1, 387 bp) and 38 iterations of repeat motif 2 (RM2, 339 bp); RM1 and RM2 predicted amino acid sequences are mostly conserved except for a few short regions (Fig. 4). These results demonstrate that previously reported AgSp1 repeat motifs were representative of a portion the C-terminal transition region, rather than the dominant repeat motifs described here (Supplementary Fig. S1, underlined sequence aligns to reported repeat of *Argiope argentata* (Fabricius, 1775) from Collin et al. (2016).

**Figure 4:**
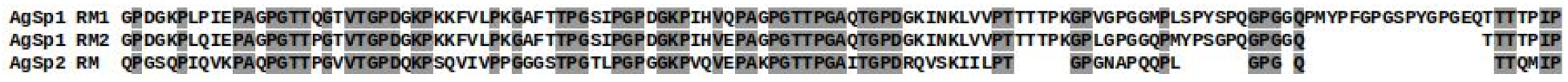
Predicted amino acid sequence of repeat motif 1 (RM1) and repeat motif 2 (RM2) from AgSp1 aligned with the repeat motif (RM) from AgSp2 of *A. trifasciata*. Conserved residues are highlighted.

In addition to *A. trifasciata*, we *de novo* assembled Illumina RNAseq data from the aggregate glands of *Argiope aurantia* Lucas (1833), and assembled publicly available aggregate gland RNAseq data sets from GenBank to discover the repeat motifs of aggregate spidroin genes from several other species: an additional orb weaver, *N. clavipes*, and three cobweb weavers, *Steatoda grossa* (C. L. Koch, 1838), *Latrodectus hesperus*, and *Latrodectus geometricus* Koch (1841) (SRR accession numbers in Supplementary Table S1). AgSp1 from a fourth cobweb species, *Parasteatoda tepidariorum* (C. L. Koch, 1841), was discovered after a BLASTp search using AgSp1 RM2 as a query (~14.8 kb; accession: XM_021148663.1). This sequence was assembled using short read data and is likely incomplete, though many repeat motifs are present and therefore included in our comparison.

Across species, repeat motifs can be classified into four subgroups with many conserved residues and completely conserved lengths, followed by a variable ‘tail’ (Fig. 5). Subgroups from all species begin with a GPXG group preceding a high proportion of hydrophobic amino acids (Fig. 5A, highlight). The first three subgroups contain regions high in serine/threonine residues, which are likely glycosylated, and are flanked by proline and glycine. The AgSp1 repeat tail region of orb weaving species is distinct from the cobweb species, which is both longer and contains stretches of poly-threonine on either end. The tail regions also contain GGQ, PGG, GPG, and QGP motifs found frequently in other spidroins, and the tail of *A. trifasciata* aligns well to repeats of major ampullate silk spidroin 2 (MaSp2; Fig. 5B, highlight). All species have several variations of their repeats, however the subgroups remain conserved with most variation again lying in the tail region (Supplementary Figure S3). Without full length sequences from the other species, it is difficult to assess if these repeat motifs are part of transitional regions or if dominating motifs are present as in *A. trifasciata*. However, the motifs chosen for comparison were the most similar to repeat motifs of the *Argiope* species.

**Figure 5:**
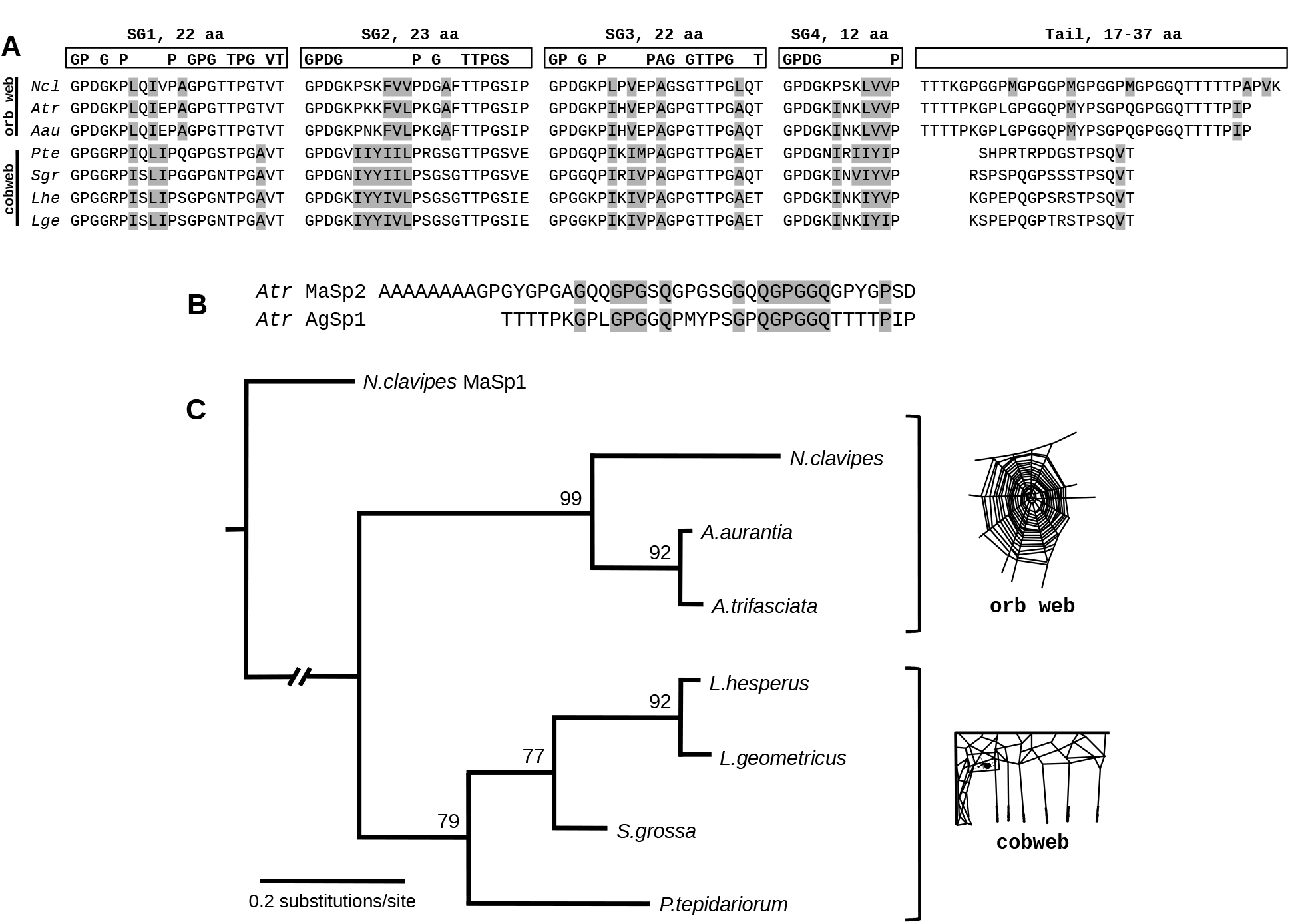
Repeat motif and phylogenetic comparisons for AgSp1 across species. (A) Repetitive AgSp1 predicted amino acid motifs for seven species, three orb web weavers and four cobweb weavers. Each repeat consists of 4 subgroups (SG) with a conserved number of amino acids (aa) followed by a variable length tail. Coneerved amino acid residues are within boxes. Grey highlight within species repeats indicates hydrophobic residues. (B) Aligned A.trifasciata major ampullate spidroin 2 (MaSp2) repeat (accesgion: AF350267.1) and AgSp1 repeat tail predicted proteins; grey highlight indicates shared amino acid residues. (C) Maximum likelihood tree with bootstrap support of repeat region encoding nucleotides (alignment Supplementary Fig. S5). Tree was rooted using the repeat region of major ampullate silk (MaSpl) from *N. clavipes* (slashed lines indicate this branch was arbitrarily shortened).

Percentages of the most prominent or glycosylation-associated amino acids that make up the AgSp1 glue repeats vary between orb web weavers and cobweb weavers (Fig. 5A, Table 2). The percent of glycine is similar between the repeats motifs of the two glue types (~22%, P = 0.7847), however orb web repeats have a higher percent of proline than cobweb repeats (23.1% vs 19.6%, respectively, P < 0.0001). The total amount of potentially O-glycosylated residues (serine + threonine) is the same between the two glue type repeats (17.8%), however orb web repeats contain more threonine than cobweb repeats (14.9% vs 10.4%, respectively, P = 0.0003) and cobweb repeats contain more serine than orb web repeats (7.3% vs 2.3%, respectively, P = 0.0006).

**Table 2:**
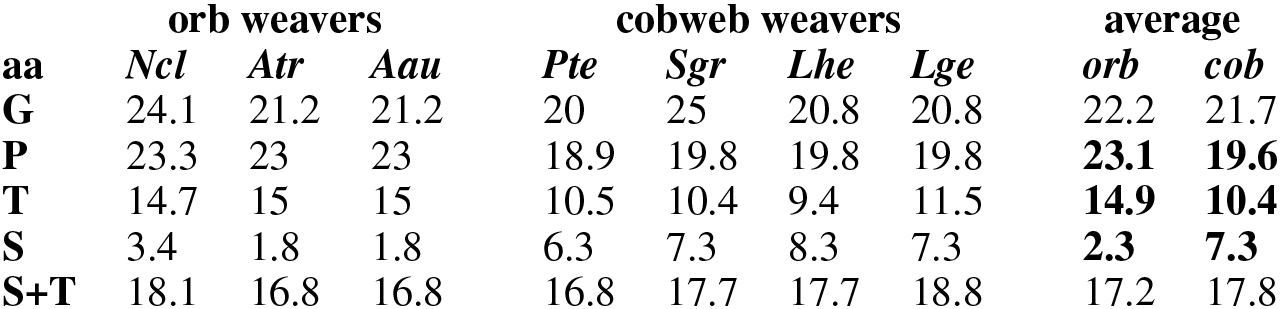
Percentage of glycosylated and glycosylation-associated amino acids (aa) in the repeat motifs of three orb web and four cobweb species. Bold indicates significant differences (*P* < 0.5).

| aa | orb weavers |  |  | cobweb weavers |  |  |  | average |  |
| --- | --- | --- | --- | --- | --- | --- | --- | --- | --- |
|  | *Ncl* | *Atr* | *Aau* | *Pte* | *Sgr* | *Lhe* | *Lge* | *orb* | *cob* |
| **G** | 24.1 | 21.2 | 21.2 | 20 | 25 | 20.8 | 20.8 | 22.2 | 21.7 |
| **P** | 23.3 | 23 | 23 | 18.9 | 19.8 | 19.8 | 19.8 | **23.1** | **19.6** |
| **T** | 14.7 | 15 | 15 | 10.5 | 10.4 | 9.4 | 11.5 | **14.9** | **10.4** |
| **S** | 3.4 | 1.8 | 1.8 | 6.3 | 7.3 | 8.3 | 7.3 | **2.3** | **7.3** |
| **S+T** | 18.1 | 16.8 | 16.8 | 16.8 | 17.7 | 17.7 | 18.8 | 17.2 | 17.8 |

We conducted maximum likelihood analyses of AgSp1 repetitive motifs for the three orb web and four cobweb weaving species rooted with MaSp1 repeats from *N. clavipes* (accession: M92913.1), a spidroin found in non-orb weaving spider groups as well (Fig. 5C). Bootstrap support for phylogenetic analysis reflects the most recent described spider relationships (Garrison et al., 2016; Bond et al., 2014; Fernández et al., 2014), as well as analyses based on the AgSp1 carboxyl-terminus (Collin et al., 2016).

### AgSp2

AgSp2 is half the size of AgSp1 and is organizationally distinct. The 5’ end is also a member of the Spidroin_N domain superfamily (BLASTx e-value 3.49e-13), however it lacks the subsequent N-terminal repeats found in AgSp1. Instead, the N-terminus leads directly into a short glutamine-rich region, and similar regions intersperse iterated AgSp2 repeat motif blocks along the length of the gene. AgSp2 has one main repeat motif (RM), which forms two main blocks of 14 and 27 iterations, as well as several smaller blocks (Fig. 3; Supplementary Fig. S2). As in AgSp1, there are repeat motifs that appear to be degenerate, and are usually located at ends of repeat blocks. The repeat motif of AgSp2 is similar to those of AgSp1 (Fig. 4, with four conserved subgroups and a unique tail. AgSp2 corresponds to AgSp-b of *N. clavipes* reported in Babb et al. (2017), however we did not find evidence for aggregate spidroins AgSp-a or AgSp-d from their report.

We identified the putative AgSp2 from cobweb weavers *L. geometricus* (accession: MK138562) and *L. hesperus* (accession: MK138563; Supplementary Fig. S6) after *de novo* assembly of online data sets (Supplementary Table S1). The assembly software was able to completely reconstruct AgSp2 in both species, with no evidence of truncation as seen in AgSp1. AgSp2 is extremely reduced to less than 2 kb in these species, and has lost the repetitious nature of most spidroins. BLASTp results identify them as flagelliform (FLAG) spidroin N-termini, however after N-terminal alignment with *A.trifasciata* AgSp2 and *A. argentata* FLAG, the encoding sequences have ~10% more bases in common with *A. trifasciata* AgSp2 (Supplementary Fig. S7).

### Gene Expression Analyses

We performed gene expression analyses using transcript data sequenced from *A. trifasciata* aggregate gland tissue to compare AgSp1 and AgSp2 expression, and from major ampullate silk gland and fat body tissues as controls. AgSp1 and AgSp2 are among the most highly expressed genes within the aggregate glands, with significantly (q < 0.05) less expression detected in the major ampullate or fat body tissues (Fig. 6A). Within the aggregate glands, expression of AgSp1 and AgSp2 is not significantly different (Fig. 6B).

**Figure 6:**
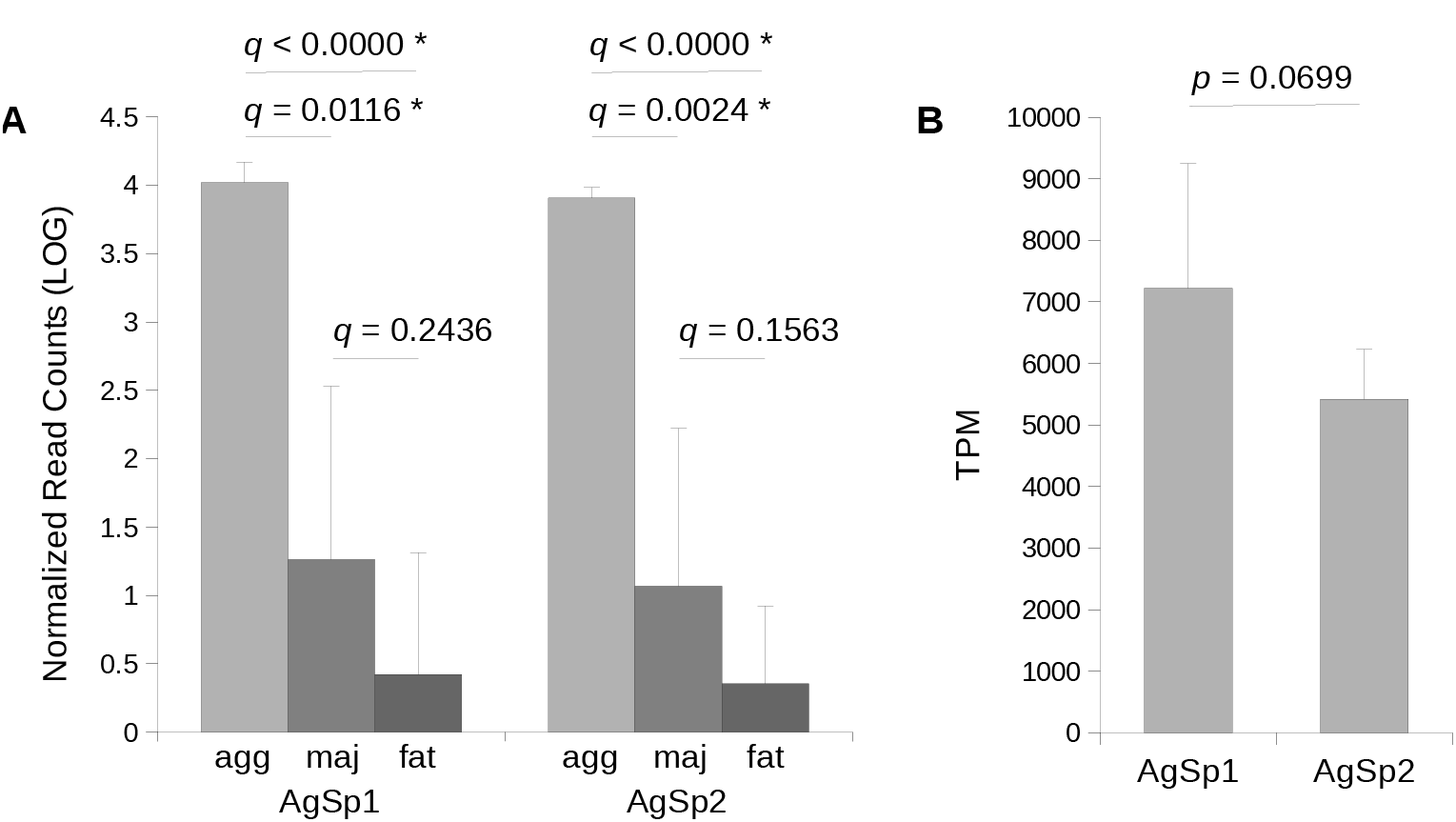
Gene expression analyses for AgSp1 and AgSp2 of *A. trifasciata*. (A) Log number of normalized reads that align to AgSp1 and AgSp2 (each manually edited to 999 bp of the 3’ ends) from aggregate gland tissue; (agg), major ampullate gland tissue (maj) and fat body tissue (fat). Line ends correspond to statistical comparisons between tissues. (B) Transcripts per million (TPM) for AgSp1 and AgSp2 averaged across six aggregate gland tissue samples. Asterisks indicate significant q- and p-values (<0.05).

## 3 Discussion

The aggregate glue spidroins AgSp1 and AgSp2 of *A. trifasciata* have coding sequences over 42 kb and 20 kb in length (respectively), both larger than the previous record-holding 19 kb aciniform prey-wrapping silk spidroin AcSp1 (Ayoub et al. (2013); Table 1). Both predicted proteins are structured as others in the family, containing repetitive central regions capped by conserved termini. The repetitive motifs are notably similar between the two spidroins, and predicted protein alignments of the repeat motifs from these two spidroins supports paralogous origins (Fig. 4). interestingly AgSp2 is highly reduced in cobweb weavers (Supplementary Fig. S6). It is unknown how AgSp1 and AgSp2 interact to form the sticky capture glue, or what properties each imparts, particularly considering the differences between AgSp2 in orb web and cobweb weavers. However we are able to make inferences about the molecular function of aggregate spidroins based on extensive research of related spidroins in the silk family.

Orb web glue is much more extensible than cobweb glue (Sahni et al., 2011a), and, interestingly, the tail region of orb web AgSp1 repeats contains a higher proportion of short motifs similar to those that contribute to the extensibility of major ampullate (MaSp) and flagelliform (FLAG) silks (Gosline et al., 1999; Hayashi et al., 1999; Adrianos et al., 2013; Malay et al., 2017). Similar to MaSp1 and FLAG, orb weaver AgSp1 repeat tails contain `GGQ` groups, and share `PGG`, `GPG`, and `QGP` groups in common with MaSp2 (Fig. 5). Furthermore, `GPG` motifs of minor ampullate spidroin proteins are significantly correlated with extensibility (Vienneau-Hathaway et al., 2017). Not surprisingly, the stretches of poly-alanine that impart silk strength to MaSp silk (Gosline et al., 1999) are lacking in the much less strong aggregate glue. The `GPG` and `QGP` regions are also found in the cobweb weaving species’ repeat motif tail regions, though the tails in these species are shorter and contain much fewer of these groups than their orb weaving counterparts (Fig. 5A, Supplementary Figure S3). If the protein backbone is responsible for imparting extensibility (how far the glue stretches) reduction in the tail region may be a response to differing functional needs for cobweb prey capture. Cobweb weavers intercept ambulatory prey that accidentally trip their gumfoot lines at low speeds. In contrast, orb weavers capture highly mobile flying or saltatory prey, and the glue may need to be more extensible in order to remain in contact with higher velocity prey items.

Notably, AgSp2 of *A. trifasciata* has 60% more total glutamine than AgSp1 despite being only half the size. A quarter of glutamine in AgSp2 is part of `QQ` motifs located in high density between blocks of iterated repeats (Supplementary Fig. S2). Similar expansion regions are also found in pyriform spidroins (Chaw et al., 2017), and `QQ` motifs have been shown to allow self-aggregation of these silk proteins into fibers (Geurts et al., 2010). AgSp2 is dramatically reduced in *Latrodectus* cobweb weavers, consisting of less than 2000 bp (Supplementary Fig. S6). Differences between orb weaving and cobweb weaving pyriform spidroins have also been noted, including a loss of the `QQ` motifs (Chaw et al., 2017).

Aggregate spider glue has been compared to the mucin family of proteins because of their glycosylated structure, viscoelastic properties (Shogren et al., 1989; Tillinghast et al., 1993; Choresh et al., 2009; Sahni et al., 2010), and highly repetitive central domains (Perez-Vilar and Hill, 1999). Our study supports similarities of aggregate spider glue to this family of secretory glycoproteins, however distinct differences can also be noted. Particularly, mucins contain cysteine-rich regions, which are joined via disulfide bonds between protein monomers resulting in the mucous net barrier or gel-like characteristics common to mucins (Desseyn, 2009; Bansil and Turner, 2006). Mucins with more cysteine residues produce stronger barriers, protecting underlying epithelial layers (Gouyer et al., 2015). It has been hypothesized that the conserved 2-3 cysteine residues in each AgSp1 terminus may form disulfide bridges between silk monomers (Collin et al., 2016), however gel-forming mucins contain cys-rich domains between the glycosylated regions as well, which are lacking in AgSp1 and AgSp2. Groups of nonpolar and hydrophobic amino acids, however, could contribute to multimerizing the individual glycoprotein monomers. Furthermore, these residues may enhance the elasticity of the glue as they pull together and exclude water (Gosline, 1978; Li et al., 2001; Bansil and Turner, 2006), which is hypothesized to contribute to elasticity in hydration-dependent flagelliform capture spiral silk (Hayashi and Lewis, 1998; Vollrath and Edmonds, 1989; Bonthrone et al., 1992). Dense hydrophobic groupings (Fig. 5A, highlight) in subgroups two and four of *A. trifasciata* AgSp1 RM2 correspond to distinct hydrophobic pockets in Kyte-Doolittle hydropathicity plots (Fig. 7). These hydrophobic residues are conserved across AgSp1 and AgSp2 repeat motifs, and both orb web and cobweb weavers. In contrast to extensibility, the glue needs to maintain elasticity, or the ability to resume its original shape, regardless of web type in order to retain prey that has contacted the threads.

**Figure 7:**
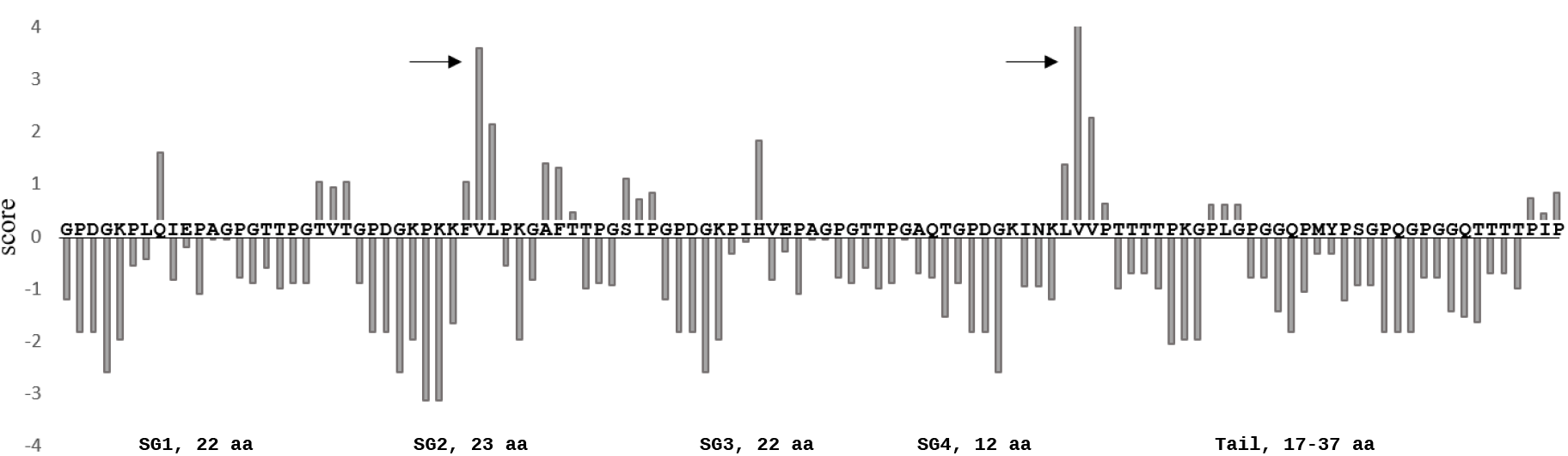
Hydrophobicity plot for repeat motif 2 of *A. trifasciata* AgSp1. Positive values indicate hydrophobicity. Arrows indicate peaks produced from triplets of hydrophobic amino acids (aa) in repeat subgroups (SG) 2 and 4 (see figure 5). Window size: 3.

Glycosylations are responsible for the adhesive qualities of spider glue (Singla et al., 2018), and at least 80% of threonine residues in Argiope glues are likely O-glycosylated with mainly N-acetylgalactosamine (GalNAc)(Dreesbach et al., 1983; Tillinghast et al., 1993). The threonine residues are also flanked by glycine and proline, which is consistent with studies that show there is an increase in these residues around O-linked glycosylated threonines (Christlet and Veluraja, 2001; Yoshida et al., 1997). Glycine is likely more prominent due to its small, non-interfering side-chains, whereas proline is hypothesized to play a structural role, kinking the protein in the form of *β*-turns. Furthermore, mucin-type glycosylated residues tend to occur in adjacent multiples, and this is consistent in the orb weaving AgSp repeats, where most threonines are directly adjacent to or within one residue from each other (Fig. 4)(Christlet and Veluraja, 2001). O-glycosylations prevent proteins from forming globules, and instead allow the protein to maintain an expanded conformation, as well as hydrating and solubilizing the protein (Shogren et al., 1989; Perez-Vilar and Hill, 1999). Electron micrographs from earlier studies of spider glue proteins showed extended, filament-like molecular structure and allowed estimations of size from 450 to 1400 kDa (figure 4 in Tillinghast et al. (1993)); our estimations from the predicted AgSp1 (1383 kDa) and AgSp2 (676 kDa) correspond to the upper and lower range values from the previous estimate.

There are differences in the percentages of amino acids that make up the AgSp1 repeats of orb webs and cobwebs (Fig. 5A). Interestingly, repeats from both glues contain a similar number of O-glycosylation sites, however orb weaver glue has a higher percentage of threonine residues than cobweb glue, while cobweb glue contains more serine residues. Changes in humidity differentially affects orb and cobweb glue adhesion (Sahni et al., 2011a), and, in addition to low molecular weight compounds and salts, this contrast in function could be due to differences between the serine and threonine glycosidic linkages (Corzana et al., 2007). These linkages structure the surrounding water in different ways, forming water pockets and bridges between the sugar and protein backbone. Additionally, threonine is more readily glycosylated than serine (O’Connell and Tabak, 1993), which could also contribute to the material differences observed between the two types of glues.

Spider silk has been notoriously difficult to synthetically produce due to the difficulty in replicating the transition from liquid protein dope to the solid fiber, which occurs within the duct of a spider’s silk gland. Aggregate spider glue is extruded as an amorphous liquid that does not undergo the same processing, however the post-translational glycosylation may provide its own challenges for future synthesis. Encouragingly, there has been success using cloned and expressed glycosyltransferases to *in vitro* O-link GalNAc residues to synthetic peptides (Yoshida et al., 1997; Hagen et al., 1997). We have identified an N-acetylgalactosaminyltransferase (pp-GalNAc-T, BLAST e-value = 3.71e-174; accession: MK138560) from *A. trifasciata*, which is part of the family of transferases that facilitates the addition of mucin-type, O-linked glycans by catalyzing the transfer of GalNAc from the sugar donor, UDP-GalNAc, to the hydroxyl groups of serine or threonine residues of the core protein, forming Ga1NAc–1-O-Ser/Thr. This specific GalNAc-T is differentially expressed in the aggregate glands compared to major ampullate gland and fat body tissues (q-value = 3.9e-8 and 7.8e-17, respectively). Furthermore, glue function does not seem to be limited by droplet size, as bolas spiders in the genus *Mastophora* produce droplets that are visibly large and specialized for capturing fast-moving moths whose wings are covered in fragile scales. *In vitro* glycosylation as well as the potential for bulk production are encouraging attributes for synthetic versions of aggregate spider glue.

## 4 Conclusion

We used Illumina short reads aligned to long reads from Oxford Nanopore’s MinlON to resolve aggregate spidroin encoding sequences AgSp1 and AgSp2 from orb weaving spider *Argiope trifasciata*. AgSp1 (42,270 bp) and AgSp2 (20,526 bp) are the largest spidroins currently described, each containing a large intron, and are of the most highly expressed genes within the aggregate glands. The predicted proteins consist of dominant central repetitive motifs and comparisons of AgSp1 repeat motifs across three orb web and four cobweb species suggests differences in the peptide backbone are related to observed functional differences of these two aggregate glue types. Aggregate spider glue must function in three ways: 1) it must adhere to surfaces, usually insect prey, 2) it must be extensible, to remain in contact with struggling prey, and 3) it must be elastic, to retain prey after extension. The tail regions of orb weaver repeats resemble regions in other spidroins that impart extensibility, while the shortened tail of cobweb weaver repeats may reflect observed reductions in this property. Conserved hydrophobic residues across the repeats of glue spidroins from both web types may contribute the elastic nature of the droplets.

As sequencing technology continues to improve, the genetics of complex biomaterials like silk spidroins will continue to be described. The similarities and differences between the orb web and cobweb glue repeat backbones are striking and provide useful insight into the relationship between form and function. There is ample opportunity to learn from differences in glue function not only between orb web and cobweb weavers, but also across the orb weavers themselves. Understanding contrasting performance optima at different humidities will provide another level of tunability to synthetic spider glues. Examining glue diversity from more distantly related species will allow an understanding of how differences in spider glue glycopeptides influence performance, contributing to the development of novel bio-inspired materials.

## 5 Methods

### Specimen collection and dissection

Adult female *A. aurantia* and *A. trifasciata* were collected from Schooley Mill Park in Highland, Maryland during September 2016 and August 2017. Specimens were kept in cages and allowed to build webs overnight. After building a fresh capture spiral in the morning, specimens were euthanized and dissected in Invitrogen RNALater (cat.no. AM7021) before 1200h. Aggregate and major ampullate glands, as well as fat body tissue samples were collected and stored in RNALater at -20°C.

### Illumina RNA sequencing and assembly

Twenty-four RNA samples were selected for Illumina sequencing: aggregate gland samples from 6 individuals of each species, and major ampullate glands and fat body tissue from 3 individuals of each species. Samples were processed according to the Qiagen RNeasy Mini Kit protocol (cat.no. 74104) including the optional on column DNAse treatment. Purified RNA was prepared for sequencing using the Illumina TruSeq Stranded mRNA library prep kit (cat.no. RS-122-2101) and run in-house on a NextSeq500 using a High Output kit with 2 x 150 PE cycles on 31 October 2016.

Sequence data sets obtained from this study and publicly available online data sets (Supplementary Table S1; http://www.ncbi.nlm.nih.gov/sra) were trimmed and filtered for quality using Trimmomatic v.0.36 (Bolger et al., 2014) and de novo assembled using Trinity (Grabherr et al., 2011) set with default parameters. Resultant contigs were searched to select any sequences that contained similar regions and motifs to aggregate spidroins previously published.

### Oxford Nanopore RNA Sequencing

Messenger RNA was extracted and pooled using the NEB Magnetic mRNA Isolation Kit (cat.no. S1550S) from 3 adult female *A. trifasciata* aggregate glands. The mRNA was then prepared for Oxford Nanopore sequencing using the Direct RNA Sequencing Kit (cat.no. SQK-RNA001). We replaced the recommended RT and buffer with those from the Thermo Scientific Maxima H Minus First Strand cDNA Synthesis kit (cat.no. K1681). The cDNA produced during library preparation is not sequenced, but provides stability for direct sequencing of the RNA molecules. Samples were run on a SpotON Flow Cell (R9.5; cat.no. FLO-MIN107), and resultant fast5 files were basecalled using Oxford Nanopore’s program Albacore v2.1.3.

### Oxford Nanopore gDNA Sequencing

Two juvenile *A. trifasciata* were collected from Schooley Mill park on July 23, 2018 and kept overnight in containers. High molecular weight DNA was gently extracted and pooled from whole, fresh specimens the next day using the MasterPure Complete DNA and RNA Purification Kit following the DNA Purification section protocol. A total of 10 μg of the gDNA extraction (pooled from both individuals) was loaded onto a Sage Science BluePippin cassette (cat.no. BLF7150) and run with a 20 kb high pass threshold overnight. The resultant elution was used directly in Oxford Nanopore’s 1D Genomic DNA by Ligation protocol (SQK-LSK109). A total of four runs were completed using SpotON Flow Cells (R9.4; cat.no. FLO-MIN106) and resultant fast5 files were basecalled using Oxford Nanopore’s program Albacore v2.2.7.

### Analyses

Sequence alignment and analysis was conducted using Geneious v11.0.5 (Kearse et al., 2012) with the Geneious alignment and ClustalW tools (Thompson et al., 1994). Maximum likelihood phylogenetic analysis was performed using the RAxML v8.2.11 Geneious plugin (Stamatakis, 2014) with the GTRGAMMA model with 10,000 bootstrap replicates. RSEM (Li and Dewey, 2011), EdgeR (Robinson et al., 2010), and R (R Core Team, 2014) were used for differential gene expression analyses. Statistical significance of amino acid percentages and TPM comparisons was calculated using Student’s t-tests. Hydrophobicity was predicted using the Kyte and Doolittle scale from online Expasy tools and a window size of 3.

**List of Abbreviations**
AgSp1: Aggregate Spidroin 1
AgSp2: Aggregate Spidroin 2
RM: Repeat Motif
NTD: N-terminal domain
NTR: N-terminal repeats
NTT: N-terminal transition
CTT: C-terminal transition
CTD: C-terminal domain
QRR: Glutamine rich region

## Funding

This work was supported by the U.S. Army Research Laboratory Postdoctoral Fellowship Program administered by the Oak Ridge Associated Universities through a contract with the U.S. Army Research Laboratory.

## Authors’ contributions

SDS contributed to experimental design, data collection, and manuscript preparation. RLR contributed to experimental design, led data collection, and reviewed and edited the manuscript.

## Acknowledgments

We thank Nathan Schwalm and Mercedes Burns for helpful comments during manuscript preparation.

---

**Figure S1:**
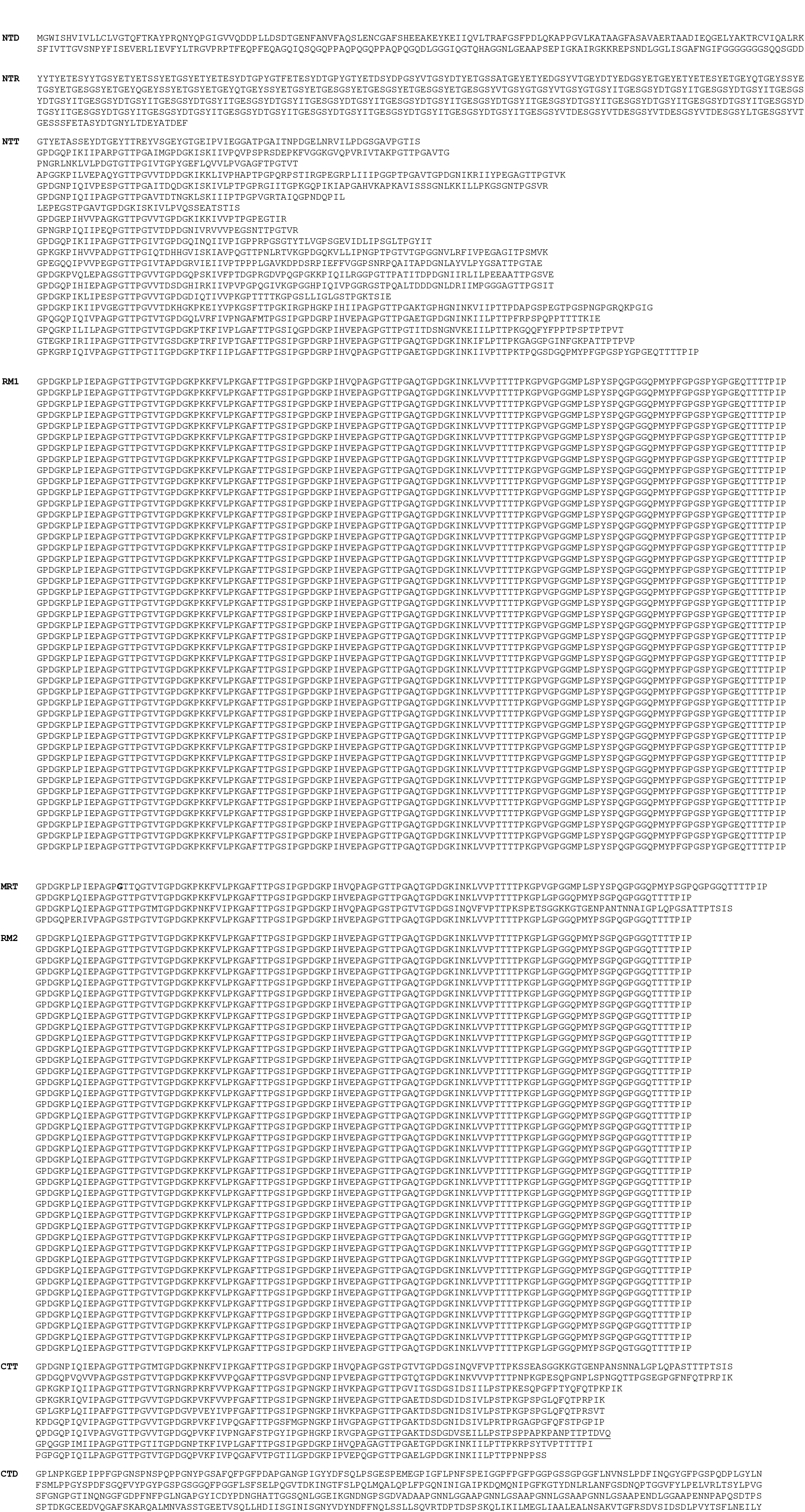
Predicted AgSp1 amino acid sequence from *A. trifasciata*. Line breaks were inserted to emphasize different regions. Abbreviations: NTD = N-terminal domain; NTR = N-terminal repeats; NTT = N-terminal transition; RM1 = repeat motif 1; MRT = mid-repeat transition; RM2 = repeat motif 2; CTT = C-terminal transition; CTD = C-terminal domain. Bolded glycine in the MRT indicates putative splice site. Underlined sequence in the C-terminal transition corresponds to the *A. argentata* repeat reported in (Collin et al., 2016). Figure is split over two pages.

**Figure S2:**
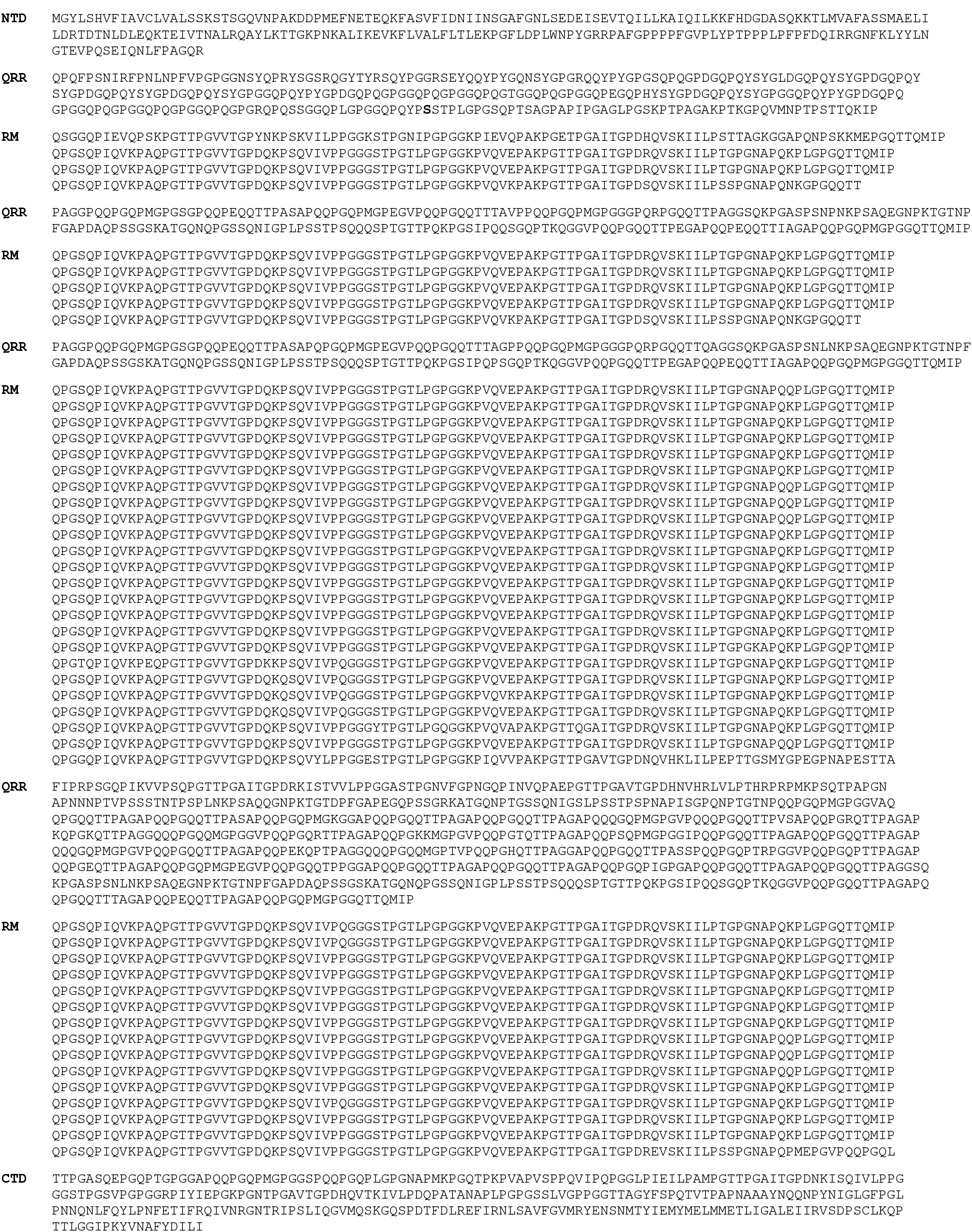
Predicted AgSp2 amino acid sequence from *A. trifasciata*. Line breaks were inserted to emphasize different regions. Abbreviations: NTD = N-terminal Domain; QRR = glutamine-rich region; RM = repeat motif; CTD = C-terminal Domain. Highlighted serine in the first QRR indicates putative splice site.

**Figure S3:**
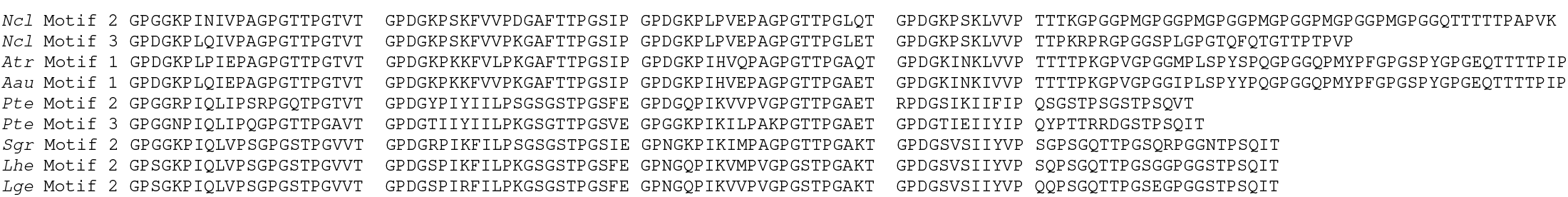
Additional AgSp1 repeat motifs from three orb weaving and three cobweb weaving species.

**Figure S4:**
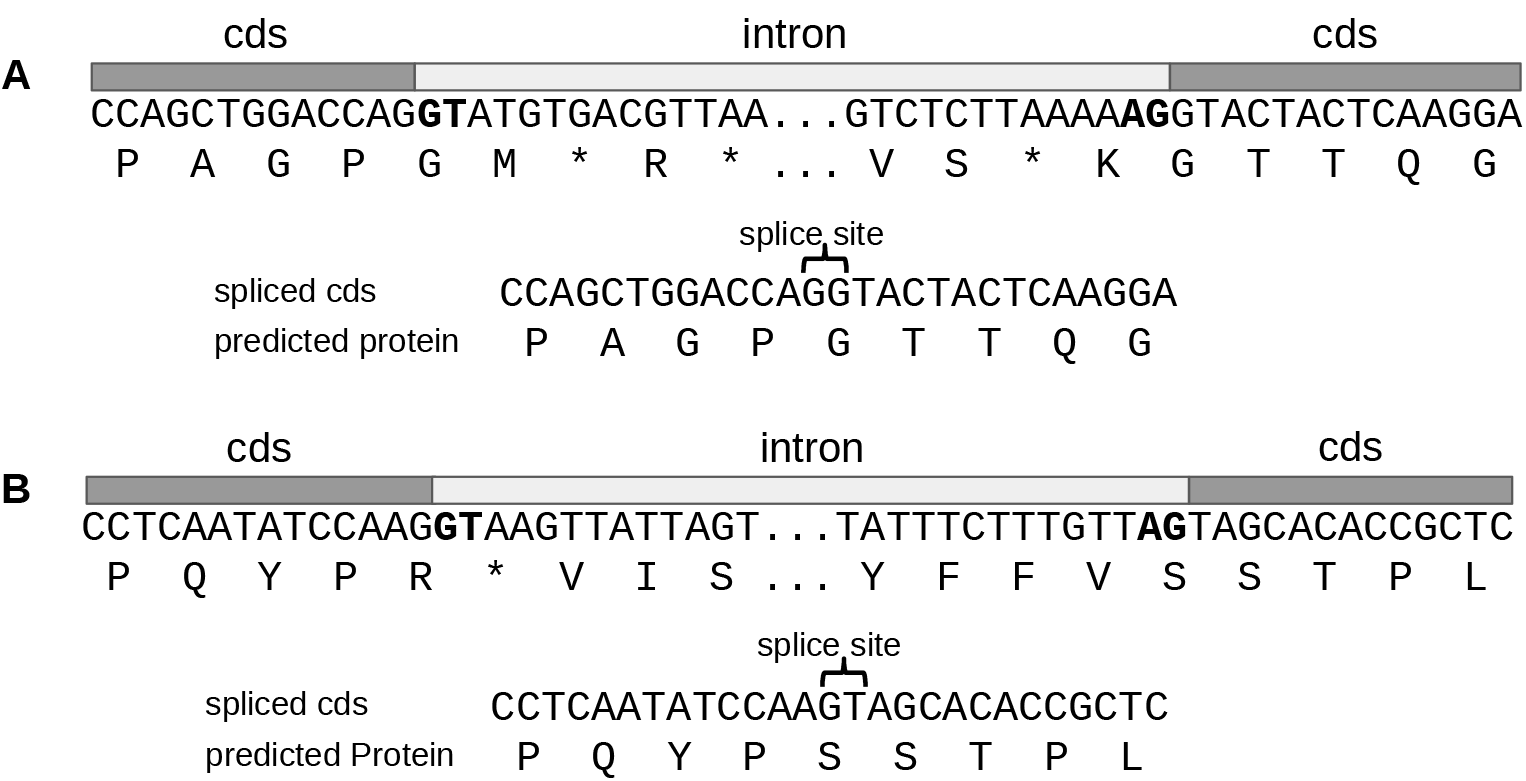
Putative splice sites for *A. trifasciata* AgSp1 (A) and AgSp2 (B) exons. Introns are capped with standard GT and AG dinucleotides (bold).

**Figure S5:**
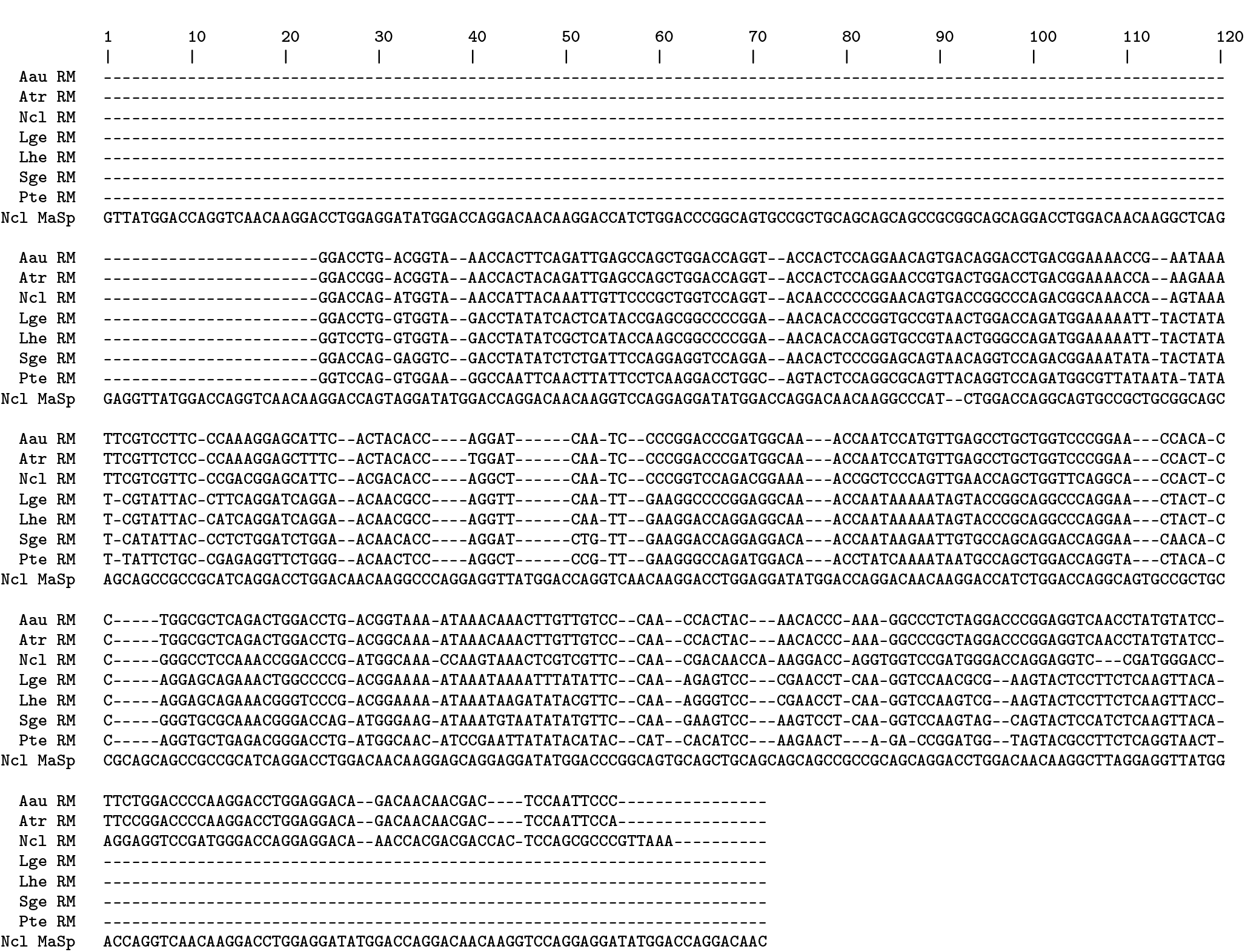
AgSp1 repeat motif nucleotide alignment from three orb weaving and four cobweb weaving spider species, and the repeat motif of major ampullate spidroin 2 (MaSp2) of orb weaver *N. clavipes*.

**Figure S6:**
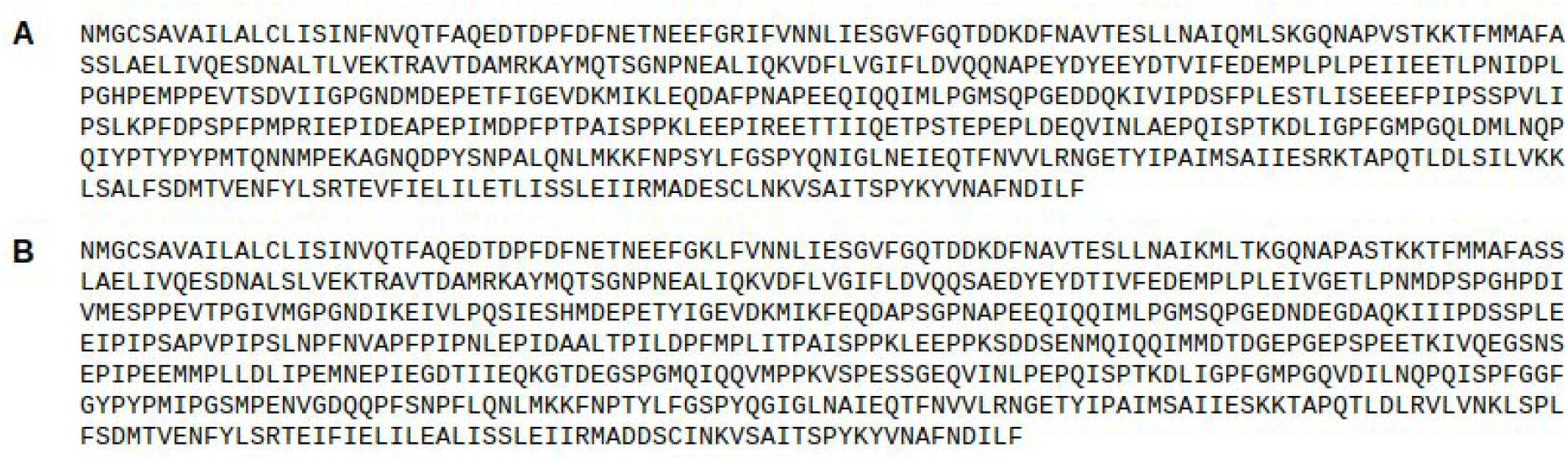
*L. geometricus* (A) and *L. hesperus* (B) AgSp2 predicted amino acid sequences.

**Figure S7:**
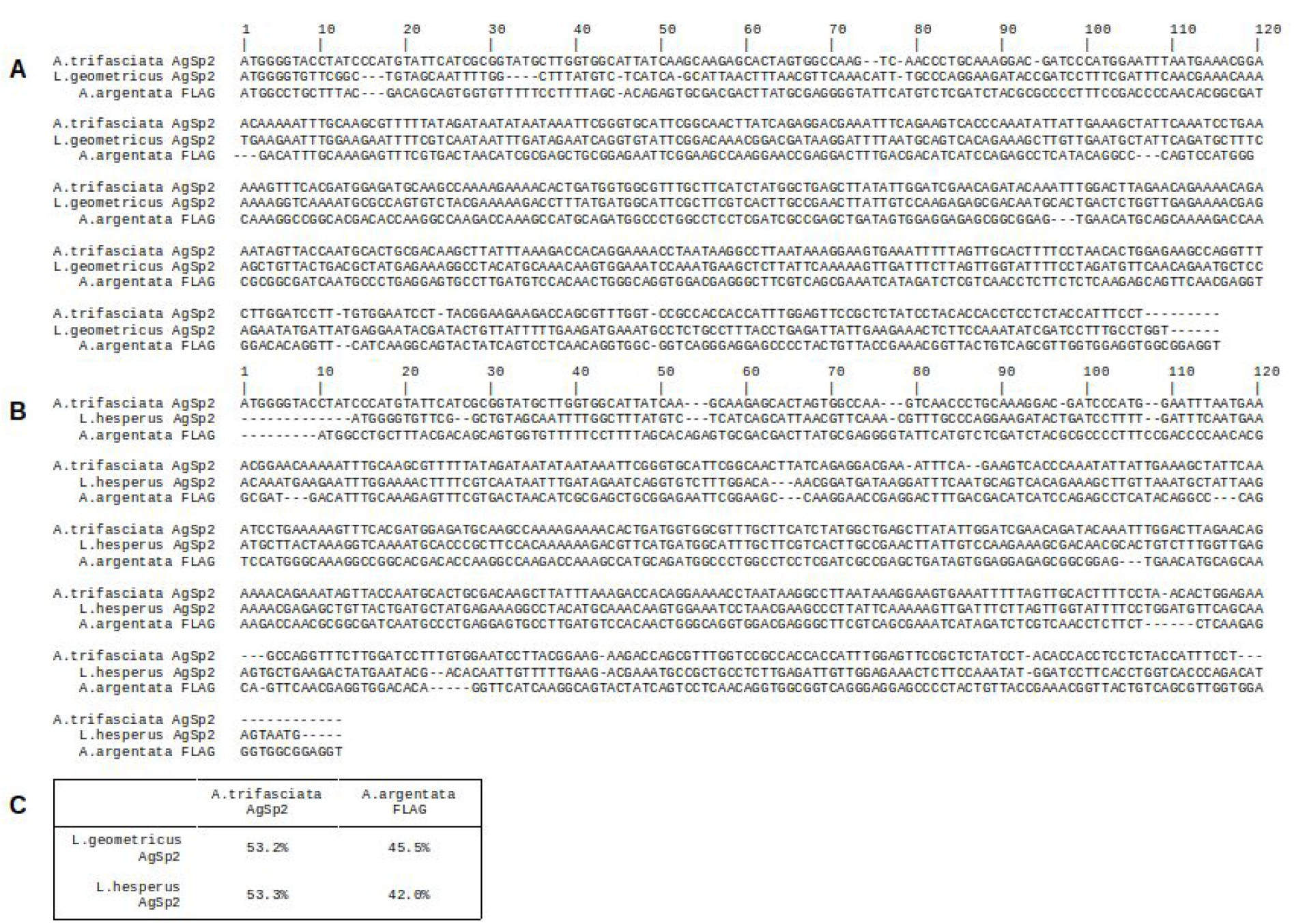
*L. geometricus* (A) and *L. hesperus* (B) AgSp2 aligned to *A. trifasciata* AgSp2 and *A. argentata* FLAG. (C) Alignment percent identity matrix.

**Table S1:**
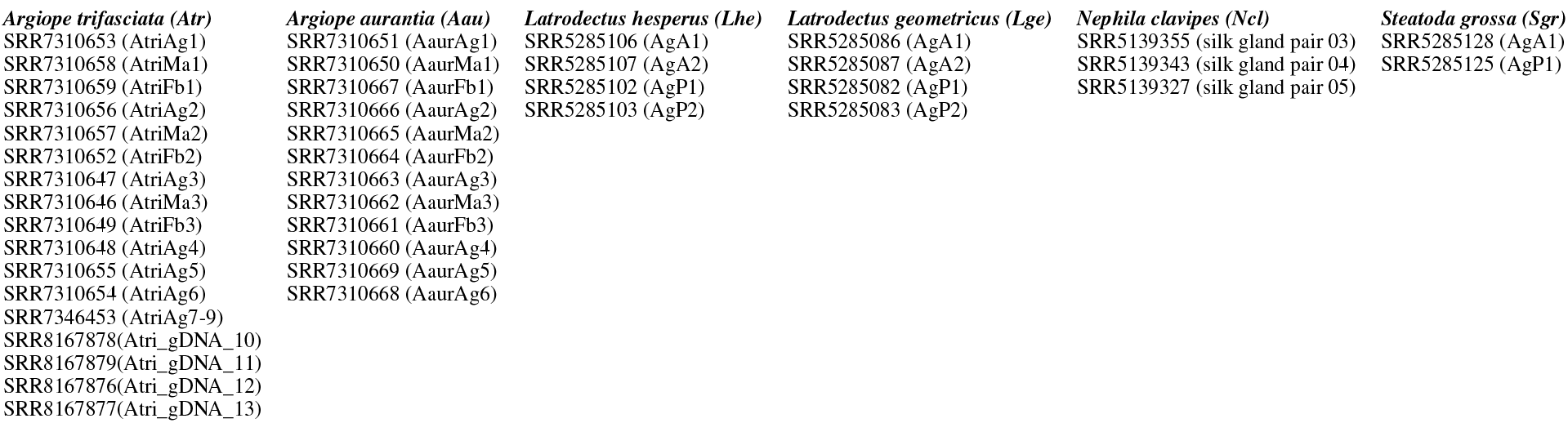
Sequencing runs used to analyze AgSp sequences. *A. aurantia* and *A. trifasciata* data were generated for this study; other species data were accessed via nCbi. All samples were sequenced using the Illumina platform except AtriAg7-13, which were sequenced using the Oxford Nanopore MinlON. Ag=aggregate, Ma=Major ampullate, Fb=Fat body, A=Anterior, P=Posterior.

